# Differential *in vitro* activity of individual drugs and bedaquiline-rifabutin combinations against actively multiplying and nutrient-starved *Mycobacterium abscessus*

**DOI:** 10.1101/2020.10.14.340422

**Authors:** Jin Lee, Nicole Ammerman, Anusha Agarwal, Maram Naji, Si-Yang Li, Eric Nuermberger

## Abstract

Current treatment options for lung disease caused by *Mycobacterium abscessus* complex infections have limited effectiveness. To maximize the use of existing antibacterials and to help inform regimen design for treatment, we assessed the *in vitro* bactericidal activity of single drugs against actively multiplying and net non-replicating *M. abscessus* populations in nutrient-rich and nutrient starvation conditions, respectively. As single drugs, bedaquiline and rifabutin exerted bactericidal activity only against nutrient-starved and actively growing *M. abscessus*, respectively. However, when combined, both bedaquiline and rifabutin were able to specifically contribute bactericidal activity at relatively low, clinically relevant concentrations against both replicating and non-replicating bacterial populations. The addition of a third drug, amikacin, further enhanced the bactericidal activity of the bedaquiline-rifabutin combination against nutrient-starved *M. abscessus*. Overall, these *in vitro* data suggest that bedaquiline-rifabutin may be a potent backbone combination to support novel treatment regimens for *M. abscessus* infections. This rich dataset of differential time-and concentration-dependent activity of drugs, alone and together, against *M. abscessus* also highlights several issues affecting interpretation and translation of *in vitro* findings.

## INTRODUCTION

In the United States and around the world, there is mounting evidence that the prevalence of chronic lung disease caused by infections with *Mycobacterium abscessus* complex has been increasing (1–3). These infections, which primarily affect vulnerable populations with underlying lung conditions such as cystic fibrosis, are particularly difficult to treat, and the limited treatment options that exist are lengthy (1–2+ years in duration), associated with severe side effects, and curative only about 50% of the time (3–5). This dearth of effective treatment options is largely due to the intrinsic resistance of *M. abscessus* complex to most available antibacterials (2); therefore, it is imperative to thoroughly investigate the potential of drugs known to have activity against this complex.

Many drugs exert differential activity depending on the status of the bacterial population, such as actively multiplying in nutrient-rich conditions versus non-replicating or slowly replicating in resource-limited environments. Among mycobacteria, this phenomenon is clearly demonstrated with the first-line tuberculosis (TB) drugs isoniazid and pyrazinamide, which have potent bactericidal activity specifically against replicating and non-replicating *Mycobacterium tuberculosis,* respectively (6). Importantly, drugs with bactericidal activity against non-replicating persister *M. tuberculosis* populations, such as pyrazinamide, bedaquiline, and rifamycins, are specifically associated with treatment-shortening in TB drug regimens (6–8). For *M. abscessus* lung disease, the role of non-replicating persisters in disease and treatment has not been established; however, the chronic nature of the disease, as well as the long duration of treatment and the frequent occurrence or relapse after treatment, indicate that bacterial persistence is likely to be an important factor.

In an effort to better understand how to maximize the use of currently available drugs for treatment of *M. abscessus* lung disease, we evaluated bactericidal activity against both actively multiplying and net non-replicating *M. abscessus* populations. Berube *et al.* and Yam *et al.* have also previously considered the importance of evaluating drug activity against non-replicating *M. abscessus*, and developed assays to generate populations of non-replicating *M. abscessus* via nutrient starvation in phosphate-buffered saline (PBS) for 4 or 6 days (9, 10). While induction of a non-replicating state by nutrient starvation cannot possibly represent all the different *in vivo* niches that might limit bacterial multiplication and induce a persister phenotype, it is one of several standard *in vitro* “persister” assays regularly used in TB drug development (6). Furthermore, as environment bacteria, *M. abscessus* complex organisms may be well-adapted to survival in nutrient-limited environments (11–13), which may contribute to the ability of these organisms to persist in *in vivo* nutrient-deprived environments such as cystic fibrosis sputum and biofilms in lung cavities and airways (14–16).

In this study, we evaluated the activity of a panel of drugs against*M. abscessus* populations actively growing in nutrient-rich broth and against those that had been nutrient-starved in PBS for up to 14 days prior to drug exposure. All drugs evaluated (amikacin, bedaquiline, clarithromycin, clofazimine, imipenem, linezolid, and rifabutin) are either currently used for treatment of *M. abscessus* lung disease and/or are considered potentially active against *M. abscessus* based on preclinical studies (4, 5, 17–22). After an initial assessment of differential bactericidal activity of each drug alone, we then evaluated the bactericidal activity of bedaquiline and rifabutin combinations, with and without a third drug, amikacin, against actively growing and nutrient-starved *M. abscessus* populations. In addition to providing a rich collection of datasets that provide important information regarding the time-and concentration-dependent activity of drugs alone and together against *M. abscessus,* this study also highlights key issues regarding the interpretation and clinical translation of *in vitro* findings.

## RESULTS

The wild type *M. abscessus* subsp. *abscessus* strain ATCC 19977 (representing a natural mixture of about 90% smooth and 10% rough colony morphotypes) was used in all experiments. On a genomic level, this commonly used laboratory strain clusters with clinical isolates within a major circulating clone group of*M. abscessus* subsp. *abscessus* (23, 24).

### *In vitro* drug activity of single drugs against actively multiplying and nutrient-starved *M. abscessus*

We first assessed the survival of *M. abscessus* in PBS plus 0.05% Tween 80; viable bacterial counts remained relatively stable for up to 83 days (nearly 12 weeks) of incubation (**Fig. S1**). We then evaluated the concentration-ranging activity of a panel of clinically relevant drugs, namely amikacin, bedaquiline, clofazimine, imipenem, linezolid, rifabutin (Fig. 1A-F, respectively; **Tables S1-S6**, respectively), and clarithromycin (Fig. 2; **Table S7**) against *M. abscessus* populations that had been nutrient-starved for 7 days (NS-7) or 14 days (NS-14) prior to drug exposure, as well as against *M. abscessus* populations actively growing in nutrient-rich cation-adjusted Mueller-Hinton broth (CAMHB). For each drug, the concentration range tested captured clinically achievable drug levels (25–31). Across these *in vitro* conditions, four general patterns of bactericidal activity were observed: (i) bactericidal in CAMHB but little or no killing in nutrient starvation conditions (linezolid, rifabutin, and clarithromycin [up to Day 7]); (ii) little or no killing in CAMHB but bactericidal against nutrient-starved *M. abscessus* (bedaquiline); (iii) bactericidal in all conditions (amikacin); and little to no killing in all conditions (clofazimine, imipenem). For all drugs tested, concentration-dependent activity was observed in permissive assay conditions. The minimum inhibitory concentration (MIC) and minimum bactericidal concentration (MBC) values for each drug and assay condition are presented in **Table S8**.

**Figure 1.**
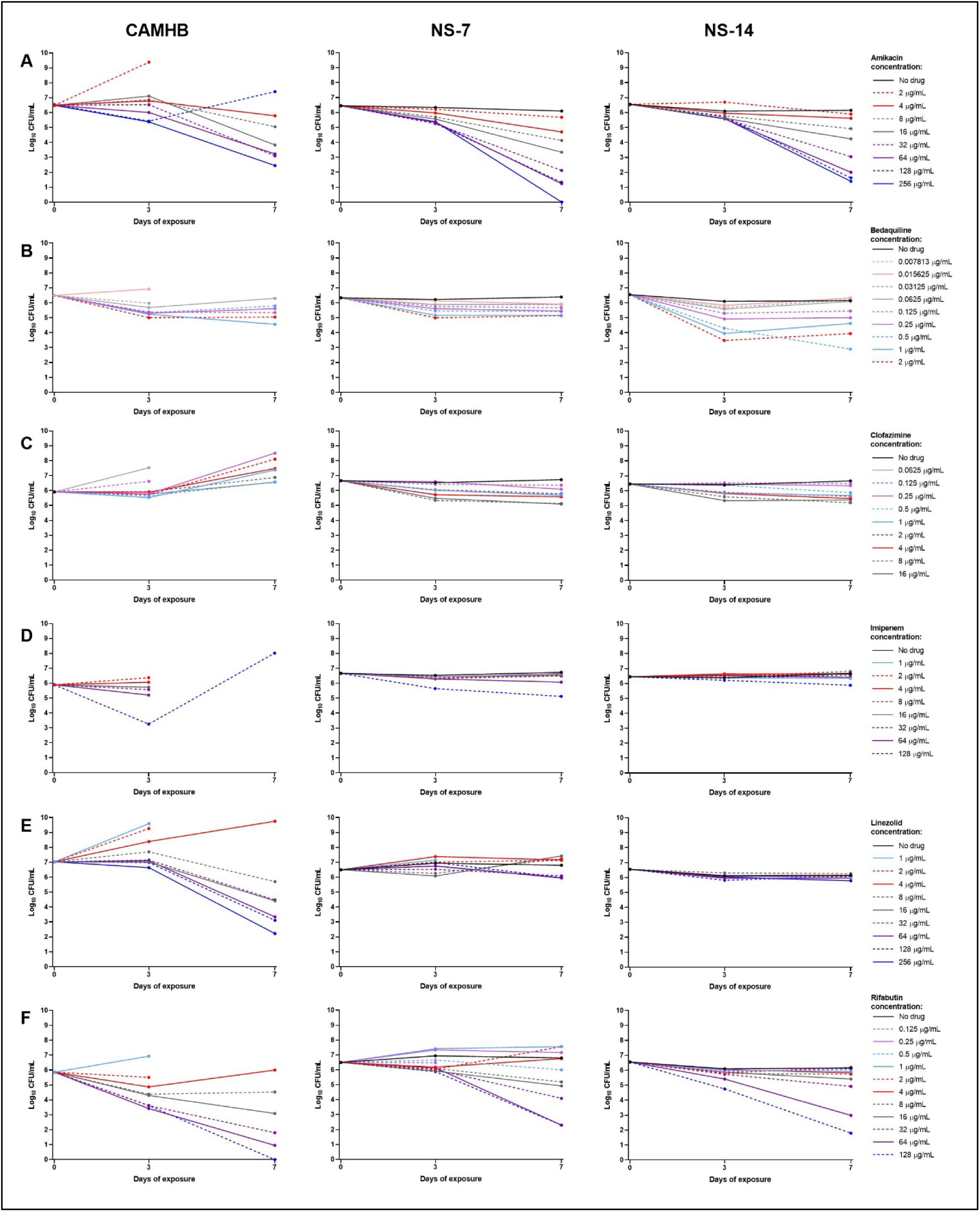
*In vitro* drug activity against actively growing and nutrient-starved *M. abscessus*. Bacterial populations were exposed to amikacin (A), bedaquiline (B), clofazimine (C), imipenem (D), linezolid (E), and rifabutin (F) for up to 7 days in the following conditions: cation-adjusted Mueller-Hinton broth (CAMHB) (left panel); nutrient-starved for 7 days prior to drug exposure (middle panel); and nutrient-starved for 14 days prior to drug exposure (right panel). Overgrowth and clumping of bacteria in CAMHB precluded CFU determination; this occurred with all no drug controls and some of the samples with lower drug concentrations at Day 3 and/or Day 7. The lower limit of detection was 0.48 log_10_ CFU/mL. All CFU data are provided in **Tables S1–S6**.

**Figure 2.**
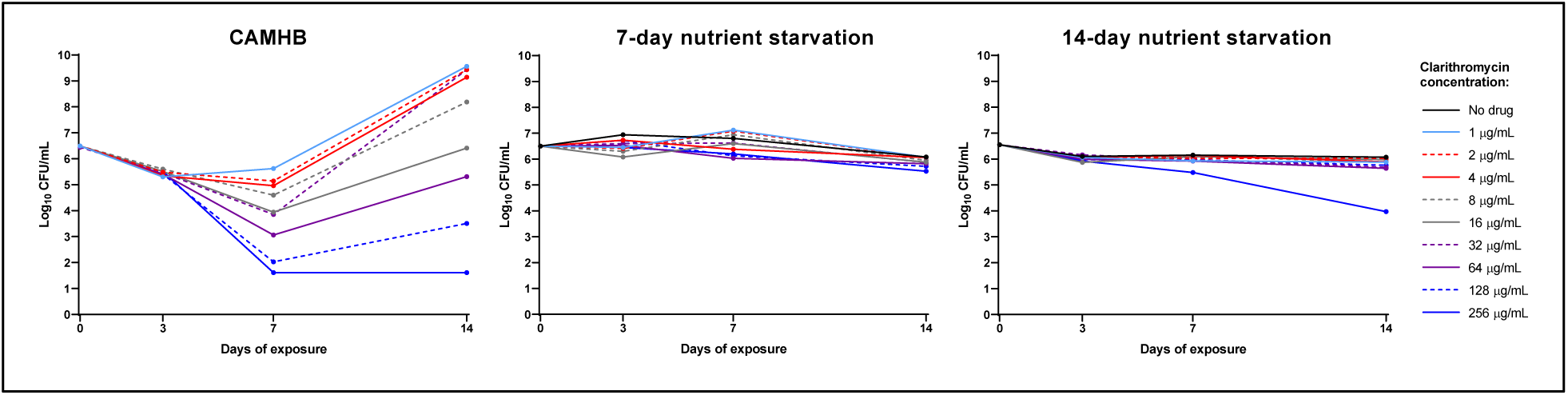
*In vitro* clarithromycin activity against actively growing and nutrient-starved *M. abscessus*. Bacterial populations were exposed to clarithromycin for up to 14 days in the following conditions: cation-adjusted Mueller-Hinton broth (CAMHB) (left panel); nutrient-starved for 7 days prior to drug exposure (middle panel); and nutrient-starved for 14 days prior to drug exposure (right panel). Over growth and clumping of bacteria precluded CFU determination in the no drug control in CAMHB were actively multiplying and overgrew/clumped, precluding accurate CFU determination. The lower limit of detection was 0.48 log_10_ CFU/mL. All CFU data are provided in **Tables S7**.

For drugs with differential activity in nutrient-rich and nutrient starvation conditions, the observed bactericidal activity against *M. abscessus* in NS-7 conditions was sometimes intermediate relative to the activity observed against bacteria in NS-14 conditions and bacteria in CAMHB. For linezolid (Fig. 1E; **Table S5**) and clarithromycin (Fig. 2; **Table S7**), activity in NS-7 was more similar to activity in NS-14 than in CAMHB. For bedaquiline (Fig. 1B; **Table S2**) and rifabutin (Fig. 1F; **Table S6**), the bactericidal activity observed in NS-7 was in between the activity observed in CAMHB and in NS-14.

Temporal differences in anti-M *abscessus* activity were also observed between drugs. Across conditions, bactericidal activity of bedaquiline was observed only during the first 3 days of drug exposure (Fig. 1B); a similar pattern was observed for the limited bactericidal activity of clofazimine in nutrient starvation conditions (Fig. 1C). Amikacin (Fig. 1A) and linezolid (Fig. 1E) had limited to no bactericidal activity during the first 3 days of exposure but were bactericidal between days 3 and 7 of exposure in nutrient starvation and CAMHB conditions, respectively. Rifabutin (Fig. 1F) had bactericidal activity during the first 3 days of exposure in CAMHB but only after Day 3 in nutrient starvation conditions. For clarithromycin (Fig. 2), concentration-independent bactericidal activity was observed during the first 3 days of drug exposure in CAMHB, and then concentration-dependent activity was observed between days 3 and 7 of exposure. Because *M. abscessus* subsp. *abscessus* is known to have inducible resistance to macrolides (32), we extended observation of clarithromycin activity up to 14 days. As expected, loss of clarithromycin activity in CAMHB was observed between days 7 and 14 of exposure.

### *In vitro* evaluation of bedaquiline and rifabutin combinations against actively multiplying and nutrient-starved *M. abscessus*

The differential activity patterns of drugs against *M. abscessus* in nutrient-rich and nutrient starvation conditions highlighted potential drug combinations which may have complementary activity in multidrug treatment regimens. Due to the growing interest in both bedaquiline and rifabutin as treatment options for *M. abscessus* infections (18, 19, 22, 33–36), we evaluated the bactericidal activity of bedaquiline and rifabutin combinations against actively growing bacteria in nutrient-rich media and non-replicating bacteria in NS-14 conditions. As it has been reported that rifabutin activity against actively multiplying *M. abscessus* can differ when assays are conducted in CAMHB or in Middlebrook 7H9 broth containing 10% (v/v) Middlebrook oleic acid-albumin-catalase-dextrose (OADC) supplement, a commonly used liquid medium for mycobacteria (17, 22), we included both types of nutrient-rich media in these assays.

First, the concentration-ranging activity of bedaquiline was evaluated alone and in combination with rifabutin at 1, 2, or 4 μg/mL. In CAMHB and 7H9 broth with 10% OADC, bedaquiline alone had limited to no bactericidal activity, although the growth inhibitory activity (on a μg/mL basis) was greater in CAMHB (Fig. 3A; **Table S2**). The MIC of bedaquiline at Day 3 increased 32-fold, from 0.0625 μg/mL in CAMHB to 2 μg/mL in 7H9 broth (**Tables S2, S8**). When combined with rifabutin, bedaquiline added concentration-dependent anti-M *abscessus* activity in both media types (Fig. 3B-D; **Table S9**). Although the activity of rifabutin alone was slightly better (on a μg/mL basis) in 7H9 broth than in CAMHB, bedaquiline added more bactericidal activity in CAMHB. For example, in the presence of rifabutin at 1 μg/mL, adding bedaquiline at 0.0625 and 0.125 μg/mL reduced Day 3 CFU counts by 1.17 and 2.54 log_10_ CFU/mL, respectively, compared to rifabutin alone in CAMHB; however, in 7H9 broth, bedaquiline at these same concentrations reduced CFU counts by only 0.58 and 0.77 log_10_ CFU/mL, respectively, and ≥2 log_10_ CFU/mL reduction compared to rifabutin alone was only achieved with bedaquiline at 2 μg/mL in 7H9 broth (Fig. 3B; **Table S9**). The magnitude of the contribution of bedaquiline decreased with increasing concentration of rifabutin. In CAMHB, the maximum bacterial reduction at Day 3 associated with bedaquiline was 2.82, 2.46, and 1.84 log_10_ CFU/mL when added to rifabutin at 1, 2, and 4 μg/mL, respectively; and in 7H9 broth, the maximum reduction associated with bedaquiline was 2.36, 1.37, and 0.61 log_10_ CFU/mL when added to rifabutin at 1, 2, and 4 μg/mL, respectively. Overall, ≥1 log_10_ CFU/mL killing between Day 0 and Day 3 was achieved at rifabutin/bedaquiline combinations of 1/0.125, 2/0.125, and 4/0.06 μg/mL in CAMHB and 1/4, 2/2, and 4/2 μg/mL in 7H9 broth. Bacterial regrowth between Day 3 and Day 7 was consistently observed in both media types.

**Figure 3.**
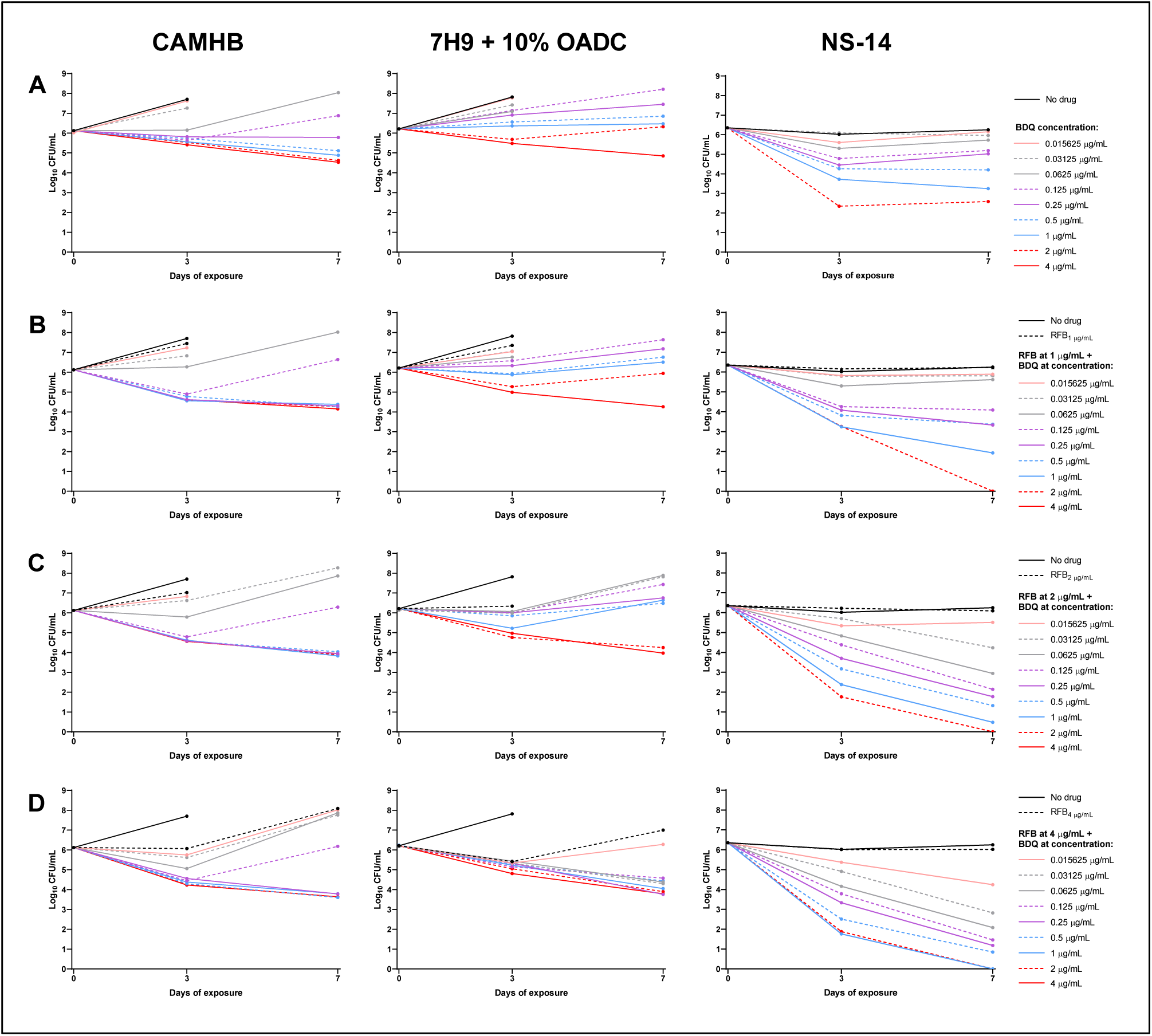
*In vitro* activity of bedaquiline (BDQ) alone and in combination with rifabutin (RFB) against actively growing and nutrient-starved *M. abscessus*. Bacterial populations were exposed to bedaquiline alone (A), or bedaquiline plus rifabutin at fixed concentrations of 1 μg/mL (B), 2 μg/mL (C), or 4 μg/mL (D) in the following conditions: cation-adjusted Mueller-Hinton broth (CAMHB, left panel); 7H9 broth with 10% Middlebrook oleic acid-albumin-dextrose-catalase (OADC) supplement (middle panel); and nutrient-starved in PBS for 14 days (NS-14) prior to drug exposure (right panel). Overgrowth and clumping of bacteria in CAMHB or 7H9 broth precluded CFU determination for the no drug controls and some samples with lower drug concentrations at Day 7. The lower limit of detection was 0.48 log_10_ CFU/mL. All CFU data are provided in **Tables S2, S9**.

As previously observed (Fig. 1B, F), bedaquiline alone had potent, concentration-dependent bactericidal activity against *M. abscessus* that had been nutrient-starved for 14 days prior to drug exposure, with all killing occurring during the first 3 days of drug exposure (Fig. 3A), and rifabutin alone at 1, 2, or 4 μg/mL had no bactericidal activity (Fig. 3B-D). During the first 3 days of exposure, the bactericidal activity of bedaquiline in the presence of rifabutin slightly increased with each escalating concentration of rifabutin, and in contrast to bedaquiline alone, this bactericidal activity extended up to Day 7 in a bedaquiline and rifabutin concentration-dependent manner (Fig. 3; **Table S9**). The rifabutin/bedaquiline concentrations that resulted in ≥2 log_10_ CFU/mL killing compared to Day 0 were 1/0.25, 2/0.25, and 4/0.125 μg/mL on Day 3 and 1/0.125, 2/0.0625, and 4/0.03125 μg/mL on Day 7.

Next, the concentration-ranging activity of rifabutin was evaluated alone and in combination with bedaquiline at 0.03125 and 0.125 μg/mL. As previously observed (Fig. 1F), rifabutin had bactericidal activity against actively multiplying *M. abscessus,* and in this set of assays, rifabutin alone had greater activity, on a μg/mL basis, in CAMHB than in 7H9 broth (Fig. 4A; **Table S6**). The MIC of rifabutin at Day 3 was similar in both media types (2 μg/mL in CAMHB and 4 μg/mL in 7H9 broth), but the concentration that resulted in ≥2 log_10_ CFU/mL killing increased at least 16-fold, from 4 μg/mL in CAMHB to >64 μg/mL in 7H9 broth (**Tables S6, S8**). In the presence of bedaquiline, rifabutin contributed bactericidal activity at lower concentrations compared to rifabutin alone. In CAMHB, rifabutin specifically contributed ≥1 log_10_ CFU/mL kill at 0.5 and 0.25 μg/mL when combined with bedaquiline at 0.03125 and 0.125 μg/mL, respectively, and in 7H9 broth, rifabutin contributed this same magnitude of killing at 2 and 4 μg/mL when combined with bedaquiline at 0.03125 and 0.125 μg/mL, respectively (Fig. 4B-C; **Table S10**). Overall, ≥2 log_10_ CFU/mL killing between Day 0 and Day 3 was achieved at bedaquiline/rifabutin combinations of 0.03125/2 and 0.125/0.25 μg/mL in CAMHB and 0.03125/16 and 0.125/8 in 7H9 broth. Bacterial regrowth between Day 3 and Day 7 was observed in both media types.

**Figure 4.**
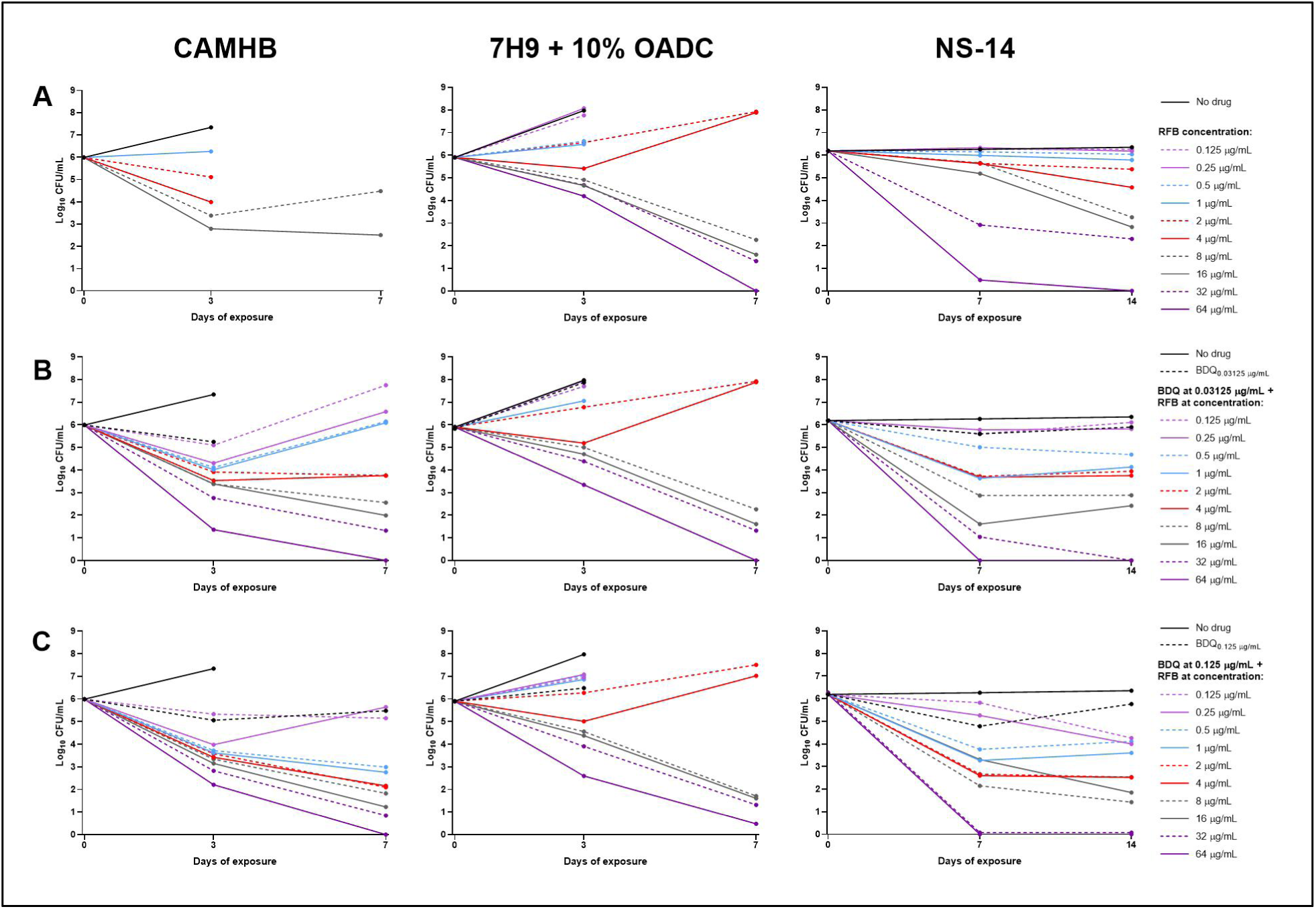
*In vitro* activity of rifabutin (RFB) alone and in combination with bedaquiline (BDQ) against actively growing and nutrient-starved *M. abscessus*. Bacterial populations were exposed to rifabutin alone (A), or rifabutin plus bedaquiline at fixed concentrations of 0.03125 μg/mL (B), or 0.125 μg/mL (C) in the following conditions: cation-adjusted Mueller-Hinton broth (CAMHB, left panel); 7H9 broth with 10% Middlebrook oleic acid-albumin-dextrose-catalase (OADC) supplement (middle panel); and nutrient-starved in PBS for 14 days (NS-14) prior to drug exposure (right panel). Overgrowth and clumping of bacteria in CAMHB or 7H9 broth precluded CFU determination for the no drug controls and some samples with lower drug concentrations at Day 3 or Day 7. The lower limit of detection was 0.48 log_10_ CFU/mL. All CFU data are provided in **Tables S6, S10**.

Because the bedaquiline concentration-ranging studies indicated that bactericidal activity of bedaquiline/rifabutin combinations in NS-14 conditions continued up to Day 7 (Fig. 3B-D), we evaluated the bactericidal activity of rifabutin concentration-ranging in combination with bedaquiline for up to 14 days of exposure. Similar to our previous result (Fig. 1F), rifabutin alone at concentrations ≤16 μg/mL had little to no activity against *M. abscessus* in NS-14 conditions (Fig. 4A; **Table S6**). However, when combined with bedaquiline, rifabutin added bactericidal activity. In the presence of bedaquiline at 0.03125 μg/mL, rifabutin specifically contributed ≥1 log_10_ CFU/mL killing at 1 and 0.5 μg/mL on Days 7 and 14, respectively, and in the presence of bedaquiline at 0.125 μg/mL, rifabutin specifically contributed ≥1 log_10_ CFU/mL killing at 0.5 and ≤0.125 μg/mL on Days 7 and 14, respectively (Fig. 4B-C; **Table S10**). Overall, the bacterial activity was similar at both concentrations of bedaquiline; the bedaquiline/rifabutin concentrations that resulted in ≥2 log_10_ CFU/mL killing compared to Day 0 were 0.03125/1 and 0.125/0.5 μg/mL on Day 7 and 0.03125/1 and 0.125/0.25 μg/mL on Day 14. In all combinations, there was little additional bactericidal activity after Day 7, and there even appeared to be some limited bacterial regrowth by Day 14.

### MIC/MBC of bedaquiline and rifabutin alone for *M. abscessus* in CAMHB and Middlebrook 7H9 broth

The two-drug combination assays plainly demonstrated that bedaquiline was less active on a μg/mL basis in 7H9 broth than in CAMHB. However, for rifabutin, the results were less clear; in one assay, rifabutin alone appeared more active in 7H9 (Fig. 3), while in another assay, rifabutin appeared more active both alone and in combination in CAMHB (Fig. 4). We therefore directly compared the MIC and MBC of each drug in both media types, using the standard reading time of 3 days. Overall, both drugs exhibited greater activity on a μg/mL basis in CAMHB than in 7H9 broth (**Fig. S2**). Based on CFU counts, the bedaquiline MIC increased 16-fold, from 0.0625 μg/mL in CAMHB to 1 μg/mL in 7H9 broth (**Fig. S2A; Table S2**). Consistent with our previous results (Figs. 1B, 3A), bedaquiline had limited to no bactericidal activity against actively multiplying bacteria in either media type. The rifabutin MIC was 4 μg/mL in CAMHB and 2 μg/mL in 7H9 broth (**Fig. S2B; Table S6**); however, the overall concentration-ranging activity of rifabutin was superior in CAMHB. As previously observed (Fig. 1F), rifabutin had bactericidal activity in CAMHB, but had much more limited bactericidal activity in 7H9 broth, a phenomenon not clearly captured by using MIC/MBC values as the read-out.

### *In vitro* evaluation of bedaquiline combined with rifabutin and amikacin against actively multiplying and nutrient-starved *M. abscessus*

In both nutrient-rich and nutrient starvation conditions, combinations of bedaquiline and rifabutin had improved bactericidal activity compared to either drug alone. In the interest of regimen-building, we next evaluated the impact of adding a third drug to the bedaquiline-rifabutin combination, and we selected amikacin due to its observed bactericidal activity against actively multiplying and nutrient-starved bacteria (Fig. 1A). The concentration-ranging activity of bedaquiline in combination with rifabutin at 1 or 2 μg/mL and amikacin at 4 or 16 μg/mL was therefore assessed against *M. abscessus* in nutrient-rich and NS-14 conditions. Due to the complexity of this study, we assessed activity in nutrient-rich conditions using CAMHB only.

Consistent with our previous findings (Figs. 1B, 3A), bedaquiline alone had little to no bactericidal activity in CAMHB (Fig. 5A; **Table S2**). When added to any concentration combination of the amikacin/rifabutin backbone, bedaquiline did not add any activity during the first 3 days of exposure; the bactericidal activity of the two-drug amikacin/rifabutin combination was not different than the activity of amikacin/rifabutin plus bedaquiline at any concentration tested (Fig. 5B-E; **Table S11**). After Day 3, concentration-dependent activity of bedaquiline was observed in CAMHB; however, bedaquiline appeared to antagonize the amikacin/rifabutin backbone when combined with the higher concentrations of amikacin and rifabutin. With backbone amikacin/rifabutin concentrations of 4/2, 16/1, and 16/2 μg/mL, the addition of bedaquiline at concentrations of <1, <1, and <0.25 μg/mL, respectively, resulted in higher log_10_ CFU/mL at Day 7 than for the two-drug backbone alone (Fig. 5C-E; **Table S11**).

**Figure 5.**
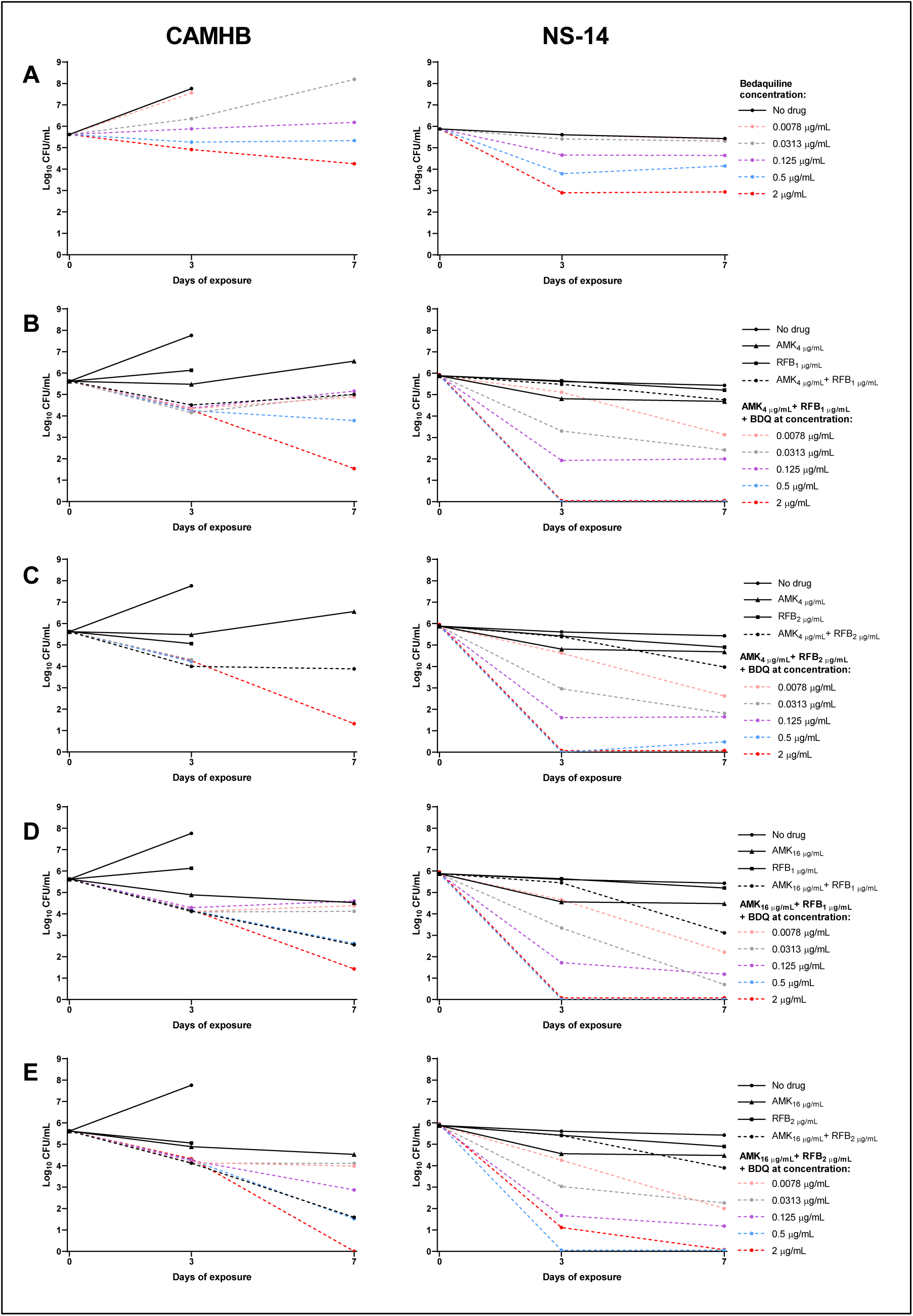
*In vitro* activity of bedaquiline (BDQ) alone and in combination with rifabutin (RFB) and amikacin (AMK) against actively growing and nutrient-starved *M. abscessus*. Bacterial populations were exposed to bedaquiline alone (A), or bedaquiline plus rifabutin at fixed concentrations of 1 μg/mL (B, D) or 2 μg/mL (C, E), and amikacin at fixed concentrations of 4 μg/mL (B, C) or 16 μg/mL (C, D), in the following conditions: cation-adjusted Mueller-Hinton broth (CAMHB, left panel); 7H9 broth with 10% Middlebrook oleic acid-albumin-dextrose-catalase (OADC) supplement (middle panel); and nutrient-starved in PBS for 14 days (NS-14) prior to drug exposure (right panel). Overgrowth and clumping of bacteria in CAMHB or 7H9 broth precluded CFU determination for the no drug controls and some samples with lower drug concentrations at Day 7. The lower limit of detection was 0.48 log_10_ CFU/mL. All CFU data, including data for additional bedaquiline concentrations not included in the graphs, are provided in Tables S2, **S11**.

As previously observed in NS-14 conditions (Figs. 1B, 3A), bedaquiline alone exhibited concentration-dependent bactericidal activity during the first 3 days of exposure (Fig. 5A; **Table S2**). In contrast to what was observed in nutrient-rich broth, the combination of bedaquiline with an amikacin/rifabutin backbone had greater bactericidal activity than the backbone alone and also greater activity than bedaquiline alone; bedaquiline added concentration-dependent bactericidal activity to each of the four backbone combinations (Fig. 5B-E; **Table S11**). In the presence of any amikacin/rifabutin combination, the bacterial killing contributed specifically by bedaquiline increased by ≥2 log_10_ CFU/mL at bedaquiline concentrations of ≤0.03125 μg/mL at Day 3 and ≤0.0078125 μg/mL at Day 7, compared to the killing attributed to bedaquiline alone. The nutrient-starved *M. abscessus* population, which started at nearly 6 log_10_ CFU/mL, fell below the limit of detection (0.48 logio CFU/mL) by Day 3 at the following concentrations of amikacin/rifabutin/bedaquiline: 4/1/0.5, 4/2/1, 16/1/0.5, and 16/2/0.5 μg/mL (**Table S11**).

## DISCUSSION

This series of extended *in vitro* studies to evaluate the anti-M *abscessus* activity of single drugs and 2-and 3-drug combinations has provided substantial data supporting several key findings, with the primary finding being that bedaquiline-rifabutin may be a promising backbone combination for building novel *M. abscessus* treatment regimens. We demonstrated repeatedly that, as a single drug, bedaquiline had bactericidal activity against nutrient-starved but not against actively growing *M. abscessus* (Figs. 1B, 3A, 5A, S2A; **Table S2**), while the opposite was observed for rifabutin (Figs. 1F, 4A, S2B; **Table S6**). Despite the differential activity of each drug alone, when combined, bedaquiline and rifabutin each contributed bactericidal activity in both nutrient-rich and nutrient starvation conditions (Figs. 3B-D, 4B-C; **Tables S9, S10**).

Rifabutin’s bactericidal activity against*M. abscessus* in nutrient-rich conditions has been previously reported (17, 22); however, the clinical relevance of this finding is complicated by interpretation of achievable exposure levels of rifabutin in patients. In adults receiving the most commonly administered daily dose of rifabutin (300 mg), the maximum plasma concentration (C max) has been reported to range from 0.38–0.53 μg/mL (37–39); however, the lung/plasma concentration ratio for a 150 mg dose was reported to range from 1.4–8.6 (37). In all *in vitro* conditions studied in here, rifabutin alone did not exhibit any static or bactericidal anti-M *abscessus* activity at concentrations <1 μg/mL (**Table S6; Table S8**). However, in the presence of bedaquiline, we have shown that rifabutin can contribute bactericidal activity at concentrations achievable in plasma. When combined with bedaquiline in CAMHB, rifabutin specifically added ≥1 log_10_ CFU/mL killing at 0.25–0.50 μg/mL, and in NS-14 conditions, rifabutin specifically contributed killing at ≤0.125 μg/mL (the lowest concentration tested) (Fig. 4C; **Table S10**). However, when combined with bedaquiline in 7H9 broth with 10% OADC, rifabutin only added bactericidal activity at concentrations ≥2 μg/mL. Sarathy *et al.* reported *in vitro* additivity between rifabutin and bedaquiline by checkerboard assay against *M. abscessus* subsp. *abscessus* strain Bamboo in 7H9 broth, with MIC readings (based on optical density) of 3.6 and 0.8 μg/mL for rifabutin alone and combined with bedaquiline, respectively; however, bactericidal activity could not be assessed using this assay method (40). Interestingly, Dick *et al.* recently reported that rifabutin alone had significant *in vivo* bactericidal activity against *M. abscessus* subsp. *abscessus* strain K21 in the lungs NOD.CB17-Prkdc^scid^/NCrCrl mice; the bacterial burden in the lungs of mice treated with rifabutin at 10 mg/kg for 10 days decreased significantly more than in the lungs of untreated mice (18). Although the pharmacokinetics (PK) of rifabutin in mice have not been well studied, mean C_max_ values in mice receiving 10 mg/kg rifabutin have been reported to range from 1.65–2.41 μg/mL (41, 42). PK issues aside, it is difficult to compare our *in vitro* findings to the *in vivo* rifabutin activity reported by Dick *et al.* because, with lung bacterial counts decreasing in untreated control mice (18), neither multiplying nor stable bacterial populations were represented in this mouse model.

That bedaquiline has bacteriostatic activity but not bactericidal activity against actively-multiplying *M. abscessus* has also been reported previously (43–45), and again the issue of clinical relevance must be addressed. In patients with MDR-TB receiving the World Health Organization-recommended bedaquiline dosing scheme (400 mg daily for the first two weeks and 200 mg thrice weekly thereafter) (46), the steady-state bedaquiline plasma concentration has been reported to be 0.9–1.2 μg/mL (47–49). Day 3 MIC values in this study for bedaquiline in CAMHB mostly ranged between 0.03125 and 0.0625 μg/mL, consistent with other reports of bedaquiline MICs against *M. abscessus* clinical isolates (19, 43, 50), while in 7H9 broth, the bedaquiline MIC (2 μg/mL) was above clinically achievable plasma concentrations (**Tables S2, S8**). When combined with rifabutin, bedaquiline specifically contributed bactericidal activity at clinically achievable plasma concentrations in all conditions tested (Fig. 3B-D; **Table S9**). Therefore, combining bedaquiline and rifabutin permitted each drug to specifically contribute bactericidal activity against actively multiplying and nutrient-starved *M. abscessus* populations at clinically relevant drug plasma concentrations.

The *in vivo* activity of bedaquiline in *M.* abscessus-infected mice has been evaluated by several groups. Lerat *et al.* reported that bedaquiline at 25 mg/kg had modest bactericidal activity against *M. abscessus* ATCC 19977 in the lungs of nude mice after 2 months of treatment (51), and Le Moigne *et al.* reported similar findings in the lungs of C3HeB/FeJ mice when bedaquiline was administered at 30 mg/kg for up to 17 days (20). In both of these studies, the bacterial burden also decreased in the lungs of untreated mice. Obregón-Henao *et al.* reported strong bactericidal activity of bedaquiline treatment (30 mg/kg for 9 days) against*M. abscessus* subsp. *abscessus* strain 103 in the lungs of GKO’’ mice, in which the bacterial counts decreased in untreated mice, and also in the lungs of SCID mice, in which the bacterial burden increased in untreated mice (21). Comparison of these *in vivo* data with our *in vitro* data is again complicated by the nature of the models, with most models experiencing natural bacterial clearance. However, for bedaquiline, a true understanding of the bactericidal activity in mice must also take into account the activity of the #-desmethyl metabolite to which bedaquiline is rapidly converted in mice (but not in humans) (52). Without knowledge of the activity of the bedaquiline #-desmethyl metabolite against *M. abscessus, in vivo* activity from mouse models cannot be directly compared to *in vitro* findings.

How bedaquiline and rifabutin may act together against *M. abscessus* is not entirely clear. That bedaquiline alone had greater anti-M. *abscessus* activity in nutrient starvation conditions is consistent with its known activity against non-replicating *M. tuberculosis in vivo* (8, 53), although it should be noted that potent bactericidal activity of bedaquiline against nutrient-starved *M. tuberculosis in vitro* has not been clearly established (6, 54). It is possible that rifabutin’s inhibitory effect on actively multiplying*M. abscessus* renders the bacteria “non-replicating” and thus more susceptible to killing by bedaquiline. However, in the presence of rifabutin at 1 μg/mL, a concentration which alone permitted bacterial growth, bedaquiline still contributed bactericidal activity (Fig. 3B) Additionally, the magnitude of the bedaquiline-specific bactericidal activity decreased with increasing concentration of rifabutin (Fig. 3B-D; **Table S9**). These data suggest the bactericidal activity of bedaquiline against actively multiplying bacteria was not solely driven by rifabutin’s net effect on bacterial growth. As a rifamycin, rifabutin inhibits DNA-dependent RNA polymerase and initiation of transcription. Several groups have reported *in vitro* synergistic activity between rifabutin and clarithromycin against *M. abscessus* (35, 36, 55), and Aziz *et al.* have specifically linked this synergism to rifabutin-induced transcriptional inhibition of genes associated with inducible macrolide resistance in*M. abscessus,* thus rendering the bacteria fully susceptible to clarithromycin (55). *In vitro* exposure of actively-growing *M. tuberculosis* to bedaquiline has been shown to induce specific transcriptional responses which may help the bacteria counteract ATP depletion, thus causing the limited or delayed activity of bedaquiline (56, 57). If bedaquiline induces a similar transcriptional response in *M. abscessus,* it is possible that co-exposure to rifabutin could inhibit this response, thus rendering the bacterial population more vulnerable to ATP depletion and killing by bedaquiline. Likewise, rifabutin may also suppress expression of genes encoding efflux pumps, such as MmpS-MmpL pump systems which are known to be involved in the efflux of and resistance to bedaquiline in*M. abscessus* (58, 59).

Similarly intriguing is the apparent bedaquiline-associated bactericidal activity of rifabutin against nutrient-starved *M. abscessus.* Typically, the rifamycins, especially rifampin and rifapentine, are considered sterilizing drugs associated with the killing of non-replicating mycobacteria *in vitro* and *in vivo* (6, 60, 61). In this study, rifabutin alone had almost no activity in NS-14 conditions but did kill nutrient-starved *M. abscessus* in the presence of bedaquiline. It is possible that additional suppression of transcription and translation caused by bedaquiline-induced ATP depletion rendered the nutrient-starved bacteria more sensitive to transcriptional inhibition by rifabutin. Clearly, additional studies are needed to better understand the nature of the combined activity of bedaquiline and rifabutin against *M. abscessus.*

A second key finding of this study was that the addition of amikacin, an inhibitor of bacterial translation, further enhanced the already potent bactericidal activity of bedaquiline and rifabutin against nutrient-starved *M. abscessus.* When combined with any concentration of amikacin and rifabutin tested, bedaquiline specifically contributed ≥1 log_10_ CFU/mL killing at extremely low concentrations of 0.0078–0.0157 μg/mL (**Table S11**), indicating that the additional suppression of protein synthesis due to amikacin further increased bacterial susceptibility to killing by bedaquiline. However, the 3-drug relationship was more complicated when applied to actively multiplying *M. abscessus.* In this setting, the rifabutin-amikacin combination was bactericidal, and not only did bedaquiline not add bactericidal activity at concentrations <2 μg/mL (**Table S11**), but also interfered with the killing of rifabutin-amikacin when bedaquiline concentrations were <0.5 μg/mL (Fig. 5D-E; **Table S11**). There are very few reported studies in which the combined activity of these drugs has been evaluated. Using the checkerboard method, Ruth *et al.* reported indifferent activity (no synergy or antagonism) between bedaquiline and amikacin in CAMHB against *M. abscessus* ATCC 19977 (45). Cheng *et al.* also used the checkerboard method and found synergy between rifabutin and amikacin for *M. abscessus* ATCC 19977 in CAMHB, but also reported that this synergy was only detected in approximately half of *M. abscessus* clinical isolates tested (35). In our study, we evaluated only two concentrations each of rifabutin and amikacin with a wide concentration range of bedaquiline, limiting our ability to understand the specific role of rifabutin and amikacin in these 3-drug combinations.

Furthermore, while the amikacin concentrations (4 or 16 μg/mL) were well below the reported plasma C_max_ values (45–85 μg/mL) in humans receiving amikacin at a standard dose of 15 mg/kg (62), the rifabutin concentrations of 1 and 2 μg/mL were just outside of the plasma concentration range at the typical human dose. Additional studies are therefore needed to specifically evaluate the role of each of these drugs in the 3-drug combination across different assay conditions.

While understanding the clinically achievable plasma concentrations of drugs is a very important consideration when designing and interpreting *in vitro* studies, this single PK parameter cannot necessarily predict drug activity *in vivo.* The impact of drugs on metabolic enzymes and transporters in the liver and gut can lead to drug-drug interactions (DDIs) which raise or lower drug levels to potentially unsafe or ineffective exposure levels. Rifamycins are known to induce CYP3A4 expression, which can lead to decreased exposures of drugs metabolized by this enzyme, including bedaquiline (30, 37, 61, 63). However, unlike rifampin and rifapentine, rifabutin is a relatively weak inducer of CYP3A4 and has not been shown to significantly impact bedaquiline exposures in humans (64, 65). As amikacin is an injectable agent primarily eliminated by the kidneys (66), its exposures are unlikely to be affected by rifabutin, and we are not aware of reported DDIs between amikacin with either rifabutin or bedaquiline.

In addition to considering the efficacy and safety implications of DDIs, we must also consider that C_max_ may not be the PK parameter driving antibacterial activity. For example, for *M. tuberculosis,* the ratio of area under the plasma concentration-time curve to MIC and the C_m_ax:MIC ratio have been shown to be important drivers of rifamycin and amikacin activity, respectively (61, 67). Therefore, having reliable MIC estimates for drugs against *M. abscessus* is critical for understanding exposure-activity relationships. This brings us to another key finding of this study, namely the differences in bedaquiline and rifabutin activity observed in CAMHB and 7H9 broth. For rifabutin, we usually observed that MIC values were similar in 7H9 and CAMHB (**Table S8**), however, we observed that the bactericidal activity of rifabutin, either alone or in combination with bedaquiline, was superior (on a μg/mL basis) in CAMHB (Figs. 3B-D, S2B; **Tables S6, S8**). Aziz *et al.* and Johansen *et al.* reported that rifabutin had lower MIC/MBC values against *M. abscessus* ATCC 19977 in 7H9 than in CAMHB (17, 22). For bedaquiline, we repeatedly observed higher MIC values in 7H9 broth compared to CAMHB (**Table S8**). Ruth *et al.* also reported that bedaquiline had a higher MIC against *M. abscessus* ATCC 19977 in 7H9 broth than in CAMHB, but also found that the median MIC of clinical isolates was lower in 7H9 than in CAMHB (45). In our work, the decreased bedaquiline activity, alone or in combination with rifabutin, in 7H9 broth, often rendered bedaquiline inactive at clinically achievable plasma levels. It is currently unclear what media condition better predicts drug activity *in vivo*, which complicates interpretation of MIC/MBC values and associated pharmacodynamic relationships.

Another key finding of this study was the differential activity of other anti-mycobacterial drugs against actively growing and nutrient-starved *M. abscessus* populations. To our knowledge, the activity of neither bedaquiline nor rifabutin against nutrient-starved *M. abscessus* has been previously reported. Berube *et al.* used a resazurin-based readout (rather than CFU counts) to evaluate the 2-day bactericidal activity of a panel of drugs against *M. abscessus* strain 103 that had been nutrient-starved in PBS with tyloxapol for 4 days prior to drug exposure (9). Similar to our findings, they found that amikacin exhibited bactericidal activity in these conditions, while clarithromycin, imipenem, and linezolid had no bactericidal activity. Yam *et al.* directly compared the 2-day MBC (≥1 log_10_ CFU/mL kill) of drugs against actively-growing *M. abscessus* strain Bamboo and against bacteria that had been nutrient-starved in PBS with tyloxapol for 6 days prior to drug exposure (10). Overall, we reported similar findings for clofazimine, imipenem, amikacin, clarithromycin, and linezolid. Although Yam *et al.* reported that clarithromycin and linezolid did not have bacterial activity in nutrient-rich conditions, CFU counts were determined after only 2 days of drug exposure. In our study, we did not see killing by each of these drugs in these conditions until between Day 3 and Day 7 of drug exposure (Figs. 1A, 1E, 2); therefore, our findings are not necessarily discrepant.

These differences in nutrient starvation models highlight an additional key finding of this work, namely that both the duration of drug exposure and the duration of nutrient starvation prior to drug exposure can significantly impact the observed activity of drugs against*M. abscessus.* For most drugs tested, the observed (or lack of) bactericidal activity did not differ in NS-7 versus NS-14 conditions, suggesting that nutrient starvation for 7 days was sufficient to observe any change in drug activity. However, for bedaquiline and rifabutin, the observed bactericidal activity in NS-7 conditions appeared intermediate to the activity in CAMHB and NS-14 conditions, indicating that bacterial populations starved for 7 or 14 days were not equivalent in terms of metabolic or other cellular programming involved in susceptibility to at least some drugs. A similar effect has been reported for the drug pyrazinamide against nutrient-starved *M. tuberculosis,* such that bactericidal activity increased with increasing duration of nutrient starvation (68). Although we cannot know which duration of nutrient starvation (if any) is most relevant to an *in vivo* condition, our data indicate that nutrient starvation for 14 days allows for better overall discrimination of drug activity between nutrient-rich and nutrient starvation conditions.

As already noted, the duration of drug exposure also impacted the assessment of drug activity. For *M. abscessus* drug susceptibility testing in CAMHB, the recommended exposure time for non-macrolides is 3 days (69). In both CAMHB and 7H9 broth, we often observed large differences in observed growth inhibition between Day 3 and Day 7, with the activity of many drugs and drug combinations decreasing after Day 3. For some drugs, this may be caused by drug instability in aqueous media, as clearly demonstrated for imipenem and other β-lactams (70, 71). Therefore, for these drugs, a read-out at 3 days of exposure may be appropriate. For other drugs, the importance and relevance of activity after 3 versus 7 days of drug exposure is less clear. For the assessment of bactericidal activity, the situation becomes even more complicated due to the diversity of time-dependent activity observed for different drugs. If, when appropriate, drug stability issues were addressed by drug supplementation, it seems that activity after 7 days, as opposed to after ≤3 days, of drug exposure could be more appropriate for understanding bactericidal activity of drugs against *M. abscessus.* In one assay, we evaluated bactericidal activity after 14 days of exposure in NS-14 conditions (Fig. 4), and we observed some apparent bacterial growth between days 7 and 14, which was unexpected in nutrient starvation conditions. Interestingly, it has been demonstrated that for other bacteria, *Escherichia coli* and *Bacillus subtilis,* multiplication can occur in nutrient starvation conditions when starved bacterial cells feed off of nutrients released from dead cells (72, 73). Although we cannot know if this phenomenon was happening in our study, this highlights that we must think critically of what an *in vitro* “nutrient-starved” environment represents. In the case of our study in NS-14 conditions, it appears that assessment of bactericidal activity was most discernable between >3 but <14 days of drug exposure. Understanding the best time point to assess activity is another critical variable in understanding concentration-activity relationships for drugs against *M. abscessus.*

Finally, the ultimate goal of this work was to provide information about the potential clinical utility of drug combinations. For*M. abscessus,* there is a notorious lack of predictability between the *in vitro* and *in vivo* activity of drugs, and several potential reasons for such discordance have been highlighted in this present work, including technical issues (media types, assay time points) as well as issues related to combining drugs. For both bacteriologic and PK factors, the activity of a drug alone does not necessarily predict the activity of a drug in combination. In TB drug and regimen development, mouse models have been pivotal for bridging the gap from *in vitro* to clinical studies (74). Ongoing efforts to develop and improve mouse models of*M. abscessus* lung infection have already shown promise (20, 21, 51, 75, 76), but nearly all are complicated by the lack of natural bacterial multiplication in the lungs. Thus, while this present series of *in vitro* studies indicates that a bedaquiline-rifabutin combination has promising activity against *M. abscessus*, further studies are needed to provide both the tools and the knowledge to translate the clinical relevance of these findings.

## METHODS

### Bacterial strain

*M. abscessus* subsp. *abscessus* strain ATCC 19977 from the American Type Culture Collection (ATCC) was used in all experiments. The colonies of this strain naturally grow in two morphotypes: smooth (about 90% of the colonies) and rough (about 10% of the colonies), and this wild type morphotype mixture was used in all assays. Aliquots of low-passage bacterial master stock were stored at −80°C.

### Drugs

Amikacin, clarithromycin, and clofazimine powders were purchased from Millipore Sigma. Imipenem and rifabutin powders were purchased from Biosynth Carbosynth, and bedaquiline and linezolid powders were provided by TB Alliance. For drug activity assays, amikacin was dissolved in distilled water; all other drugs were dissolved in dimethyl sulfoxide (DMSO). Drug solutions were filter-sterilized prior to use.

### Media

The standard liquid growth medium used to initiate all bacterial cultures was Middlebrook 7H9 broth supplemented with 10% (v/v) Middlebrook OADC supplement, 0.1% (v/v) glycerol, and 0.05% (v/v) Tween 80. For drug activity assays, two types of media were used: Middlebrook 7H9 broth supplemented with 10% (v/v) OADC and 0.1% (v/v) glycerol and CAMHB; note that Tween 80 was not included in the media when drug activity was assessed. The standard solid growth medium used for determination of CFU counts was non-selective 7H11 agar supplemented with 10% (v/v) OADC and 0.1% (v/v) glycerol (referred to as “non-selective 7H11 agar”). Agar plates contained 20 mL agar in 100 x 15 mm disposable polystyrene petri dishes. Difco BBL Mueller Hinton II broth (cation-adjusted) powder (*i.e.*, CAMHB powder), Difco Middlebrook 7H9 broth powder, Difco Mycobacteria 7H11 agar powder, and BBL Middlebrook OADC enrichment were manufactured by Becton, Dickinson and Company. Glycerol and Tween 80 were purchased from Fisher Scientific.

### Drug activity assays in nutrient-rich media

To start each experiment, a stock vial of *M. abscessus* (frozen in standard growth medium) was thawed, added to fresh growth media, and incubated at 37°C, shaking, until the optical density at 600 nm (OD_600_) of the bacterial suspension reached around 1 (approximately 10^7^–10^8^ CFU/mL), at which point assays were initiated (*i.e.*, this was Day 0). Cultures were then diluted with appropriate assay media to an OD_600_ of 0.1. All drug activity assays were performed by broth macrodilution in a total volume of 2.5 mL in 14-mL round-bottom polystyrene tubes with screw caps. Two-fold dilutions of drug stock solutions were added to appropriate assay media in a total volume of 2.4 mL per tube, with the concentration of DMSO never exceeding 4% (v/v). Then, 0.1 mL of the prepared bacterial suspension (diluted to OD_600_ of 0.1) was added to each tube of drug-containing media. Tubes were vortexed and incubated at 37°C without shaking for the duration of the study. The Day 0 inoculum, as well as samples at each time point, were cultured for CFU determination.

### Bacterial survival and drug activity assays in nutrient starvation conditions

*M. abscessus* stock was cultured in growth media to an OD_600_ of around 1 as described for assays in nutrient-rich media. The bacterial suspensions were then washed 3 times in PBS with 0.05% Tween 80 as follows: culture was spun at 1900 rcf for 10 minutes, supernatant was removed, and cells were resuspended to the original volume in PBS with 0.05% Tween 80. After the third wash, the resuspended bacteria were incubated for nutrient starvation at 37°C for the indicated duration. To monitor bacterial survival, samples were removed and cultured for CFU determination at the indicated time points. For drug activity assays, after the appropriate duration of nutrient starvation (7 or 14 days), the bacterial suspension was diluted with PBS to an OD_600_ of 0.1 on Day 0 of the assay. Two-fold dilutions of stock drug solutions were added to PBS without Tween 80 in a total volume of 2.4 mL, and assay tubes were otherwise prepared as described for drug activity assays in nutrient-rich media, except that PBS without Tween was used in place of media.

### Quantitative cultures and CFU counting and analysis

In all experiments at all time points, CFU determination was done by plating serial 10-fold dilutions of bacterial suspensions on non-selective 7H11 agar plates. CFU determination was done for all samples except when the bacteria had overgrown and fallen out of suspension in a large clump that could not be visually dispersed by vortexing. Serial dilutions were made in PBS without Tween by adding 0.1 mL bacterial suspension (from assay tube or from previous 10-fold dilution) to 0.9 mL PBS.

Undiluted up to the 10^−5^ dilution were prepared for most samples, although the dilution range was extended for some samples when a higher bacterial burden was anticipated. For plating, 0.5 mL was spread across the surface of the agar. Once all liquid was absorbed into the agar, the plates were sealed in plastic bags and incubated at 37°C for 5–7 days. CFUs were then counted/recorded for each plate. The dilution that yielded CFU counts between 10–120 and closest to 50 was used to determine CFU/mL. The CFU/mL value (x) was log transformed as log_10_ (x + 1) prior to analysis.

### MIC and MBC definitions

The visual MIC was defined as the lowest drug concentration that inhibited any bacterial growth as observed by the naked eye. The MIC based on CFU counts was defined as the lowest drug concentration that inhibited growth by ≤0.1 log_10_ CFU/mL compared to Day 0. The MBC (≥1 log_10_ kill) and MBC (≥2 log_10_ kill) were defined as the lowest concentration that decreased the bacterial count by >1 or >2 log_10_ CFU/mL, respectively, compared to Day 0.

## ACKNOWLEDGEMENTS

This work was funded by the National Institutes of Health (R21-AI137814 to EN) and by the Cystic Fibrosis Foundation (EN).

